# Adrenergic signaling gates astrocyte responsiveness to neurotransmitters and control of neuronal activity

**DOI:** 10.1101/2024.09.23.614537

**Authors:** Kevin A. Guttenplan, Isa Maxwell, Erin Santos, Luke A. Borchardt, Ernesto Manzo, Leire Abalde-Atristain, Rachel D Kim, Marc R. Freeman

**Affiliations:** Vollum Institute, Oregon Health and Sciences University; Portland, Oregon, USA; Neuroscience Institute, NYU Grossman School of Medicine; New York, NY., USA

## Abstract

How astrocytes regulate neuronal circuits is a fundamental, unsolved question in neurobiology. Nevertheless, few studies have explored the rules that govern when astrocytes respond to different neurotransmitters *in vivo* and how they affect downstream circuit modulation. Here, we report an unexpected mechanism in *Drosophila* by which G-protein coupled adrenergic signaling in astrocytes can control, or “gate,” their ability to respond to other neurotransmitters. Further, we show that manipulating this pathway potently regulates neuronal circuit activity and animal behavior. Finally, we demonstrate that this gating mechanism is conserved in mammalian astrocytes, arguing it is an ancient feature of astrocyte circuit function. Our work establishes a new mechanism by which astrocytes dynamically respond to and modulate neuronal activity in different brain regions and in different behavioral states.

## Main Text

Astrocytes causally regulate neuronal circuits (*1*). Not only do astrocytes display complex calcium transients in response to neuronal activity, but astrocyte calcium influx is necessary and sufficient to regulate the firing of specific circuits with reliable behavioral consequences (*2, 3*). This represents an understudied layer of neuronal circuit modulation that must be understood to decipher how the brain encodes information. Nevertheless, surprisingly little is known about precisely what signals astrocytes respond to *in vivo*, the kinds of computations they perform, the mechanisms by which they regulate neuronal activity, and how these features may vary across brain regions or behavioral states (*4*).

Complicating this understanding, astrocyte intracellular signaling pathways behave in fundamentally different ways than those of neurons. For instance, neurons typically increase calcium activity in response to G_⍺q_ signaling and decrease activity in response to G_⍺i_ signaling (*5*). In contrast, astrocytes increase calcium activity in response to both (*6-9*). Further, while most calcium activity in neurons can be understood to represent action potential generation and the calcium-dependent release of neurotransmitters, distinct methods of increasing calcium activity in astrocytes lead to vastly different effects on the activity of downstream neuronal circuits (*6-8*). Finally, astrocytes exhibit highly varied responses to the same neurotransmitters across brain regions, times, and behavioral states of animals (*10-17*). Some studies have observed that a single neurotransmitter release event can simultaneously increase and decrease calcium levels in neighboring astrocytes within the same organism (*13*). Together, these studies emphasize that, to understand the role that astrocytes play in neuronal circuits and behavior, we must gain a better understanding of the mechanisms that govern their responses to neurotransmission.

## Results

### Tyramine gates the response of astrocytes to other neurotransmitters

To understand the molecular mechanisms that determine how astrocytes respond to different neurotransmitters, we utilized astrocytes of the larval *Drosophila* ventral nerve cord (VNC), the equivalent of the mammalian spinal cord. *Drosophila* VNC astrocytes have been shown to share the vast majority of known functions of mammalian astrocytes (*18*) and were previously used to identify specific molecules that astrocytes use to causally regulate intrinsic calcium signaling and downstream neuronal activity (*2*), a mechanism subsequently validated in vertebrates (*3*). Employing an established method to image astrocytes in intact larval brains *ex vivo*, we bath applied neurotransmitters in the presence of tetrodotoxin (TTX) to block neuronal activity and assessed direct astrocyte responses (Fig. 1A, astrocytes manipulated using the *alrm-Gal4* driver). In this context, astrocytes reliably responded to octopamine and tyramine - the functional homologues of epinephrine/norepinephrine – with whole-cell calcium responses across the entire VNC, but were unresponsive to all other neurotransmitters tested (Fig. 1B-D, S1A-B) (*2*). Given the evidence that animal behavior state and adrenergic signaling can change astrocyte calcium activity in response to neuronal firing (*19, 20*), we next asked if pre-exposure of astrocytes to tyramine could change their response to other neurotransmitters. Remarkably, approximately three minutes after exposure to tyramine (with or without tyramine washout), astrocytes became capable of exhibiting robust responses to dopamine, glutamate, acetylcholine, and GABA (Fig. 1E-F, S1C-E). This indicates that pre-exposure to tyramine can control the response of astrocytes to other neurotransmitters in the larval VNC, which we term “gating.” This response was specific to tyraminergic signaling – glutamate and acetylcholine could not elicit a response following dopamine, suggesting that some aspect of the tyramine signaling cascade was necessary for gating (Fig. 1F).

**Fig 1:**
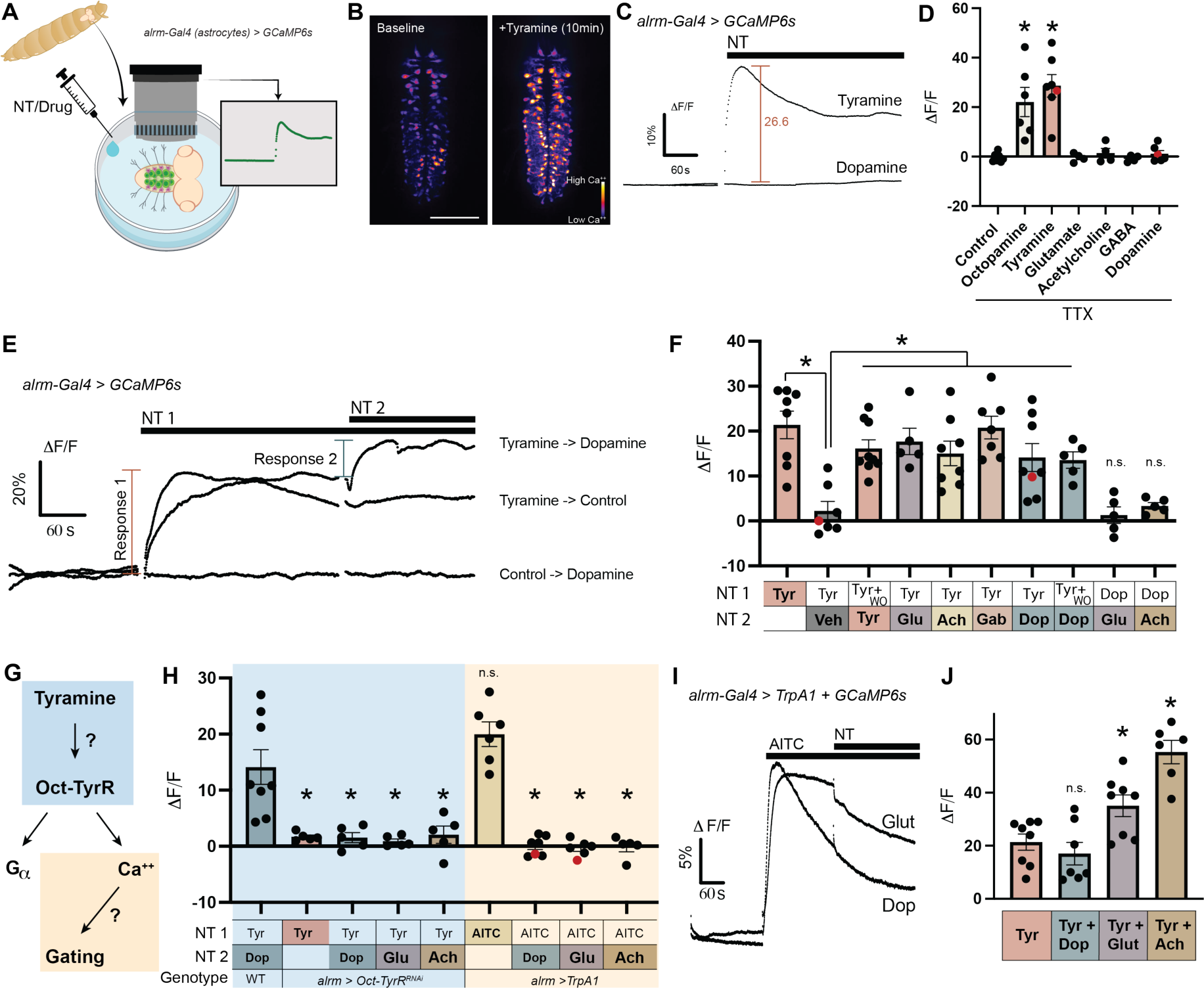
Tyramine gates the response to other neurotransmitters via the Oct-TyrR. (**A**) Schematic demonstrating the *ex vivo* preparation used to image larval ventral nerve cord (VNC) astrocytes in an intact nervous system. (**B**) VNC astrocytes before and after exposure to tyramine with GCaMP6s fluorescence pseudo-colored to demonstrate the increase in calcium level across nearly all astrocytes in response to tyramine. (Scale bar = 100 μm) **(C)** Example trace of a field of VNC astrocytes responding to tyramine with a calcium influx but showing no response to dopamine as well as demonstration of the quantification method for the tyramine response. (**D**) Quantification of astrocyte response to various neurotransmitters (NT). Note that astrocytes respond robustly to octopamine and tyramine but not to other neurotransmitters. (**E**) Example traces of astrocytes responding to tyramine followed by a second neurotransmitter (or vehicle control, Veh). Note that astrocytes respond to dopamine after tyramine stimulation but not to dopamine alone. (**F**) Quantification of astrocyte calcium responses shows that astrocytes respond to all tested neurotransmitters following tyramine exposure but not following dopamine exposure. We term this response to neurotransmitters after tyramine exposure “gating.” (WO = washout; Ach = acetylcholine, Glu = glutamate, Tyr = tyramine, Gab = GABA) (**G**) Schematic demonstrating a hypothesized pathway of gating via Oct-TyrR. (**H**) Quantification of astrocyte calcium responses showing that the gating of all tested neurotransmitters is dependent on Oct-TyrR but not dependent on calcium influx via the TRP channel TrpA1. This suggests that gating does not occur due to calcium influx similar to that induced by Oct-TyrR activation. (**I**) Example traces of *TrpA1*-expressing astrocytes responding to AITC exposure but not to subsequent glutamate or dopamine exposure. (**J**) Adding tyramine at the same time as dopamine (Tyr + Dop) does not lead to a larger calcium response than tyramine alone. In contrast, adding glutamate or acetylcholine with tyramine leads to a significantly larger calcium response than tyramine alone.* Indicates p-value < 0.05; details of statistical comparisons and exact p-values in Table S1. All error bars represent SEM. Red dots in bar graphs correspond to traces chosen as example.

We previously found that whole-cell astrocyte calcium responses to tyramine depend on a single G-protein coupled receptor (GPCR), the Octopamine-Tyramine receptor (Oct-TyrR) (Fig. 1G, blue field) (*2*). Knocking down *Oct-TyrR* specifically in astrocytes abolished both the tyramine response and the gating response to all neurotransmitters tested, suggesting that the gating mechanism is dependent on signaling downstream of Oct-TyrR (Fig. 1H). Octopamine, which can also stimulate the Oct-TyrR, could similarly elicit gating (Fig S1F). One hypothesis for how tyramine stimulation could gate a secondary response is via the calcium influx in astrocytes, which might fill the calcium reserves necessary for a secondary response or mediate a calcium-dependent intracellular signal that allows gating (Fig. 1G, orange field). To test this hypothesis, we expressed *TrpA1* in astrocytes, as this channel passes calcium in response to the exogenous ligand allyl-isothiocyanate (AITC) and has been shown to functionally replace endogenous Oct-TyrR-dependent calcium signaling in the astrocytic regulation of neuronal circuits (*2*). Stimulating calcium influx via TrpA1 did not facilitate gating of any neurotransmitters despite inducing a similar calcium response magnitude as tyramine (Fig. 1H-I). Thus, while TRP channel-mediated calcium entry itself drives astrocyte-mediated changes in neural circuit activity and animal behavior (*2*), it is not sufficient to mediate gating of astrocyte responses to glutamate, acetylcholine, or dopamine.

To better understand how tyramine might be gating other neurotransmitter responses, we added tyramine simultaneously with dopamine, glutamate, or acetylcholine. While simultaneous addition of tyramine and dopamine did not lead to a response larger than that triggered by tyramine alone, both glutamate and acetylcholine led to significantly larger combined responses (Fig. 1J). This shows that, while all gated responses required the Oct-TyrR, the kinetics of dopamine vs glutamate/acetylcholine gating appear to differ. In the case of dopamine, the lack of a simultaneous enhancement from tyramine and dopamine together implies that astrocytes undergo some form of intracellular signaling to facilitate dopamine responsiveness, a finding supported by a more in-depth analysis of the time course of dopamine gating (Fig. S1G).

### Dopamine gating is dependent on G_⍺i_ GPCR regulation of Dop2R internalization

We next asked if the GPCR signaling downstream of Oct-TyrR is responsible for gating (Fig. 2A). Expressing a cAMP indicator in astrocytes, we found that tyramine exposure led to a decrease in cAMP levels, which aligns with previous reports that, in cell lines, Oct-TyrR functions as a G_⍺i_-coupled GPCR (Fig. 2B) (*21*). To validate that G_⍺i_ signaling itself is necessary for gating, we overexpressed the pertussis toxin alpha subunit (*PTXa*), which inhibits G_⍺i_ signaling, and separately knocked down the G_⍺i_ protein in astrocytes. In both instances, disrupting G_⍺i_ signaling prevented gating of dopamine, glutamate, and acetylcholine, confirming that G-protein signaling downstream of Oct-TyrR is required (Fig. 2C). Because G_⍺i_ signaling leads to a decrease in cAMP levels, we next tested whether modulating cAMP levels without Oct-TyrR stimulation could trigger the astrocyte gating response. Indeed, pre-treatment of astrocytes with the adenylyl cyclase inhibitor SQ22536 - but not the adenylyl cyclase activating drug forskolin - was sufficient to gate the astrocyte response to dopamine (Fig. 2C-D). Interestingly, cAMP modulation was not sufficient to gate the responses to glutamate or acetylcholine (Fig. 2C). Thus, in line with the different kinetics of dopamine and glutamate/acetylcholine gating (Fig. 1J), the mechanisms of gating of neurotransmitters diverge at the level of cAMP regulation.

**Fig 2:**
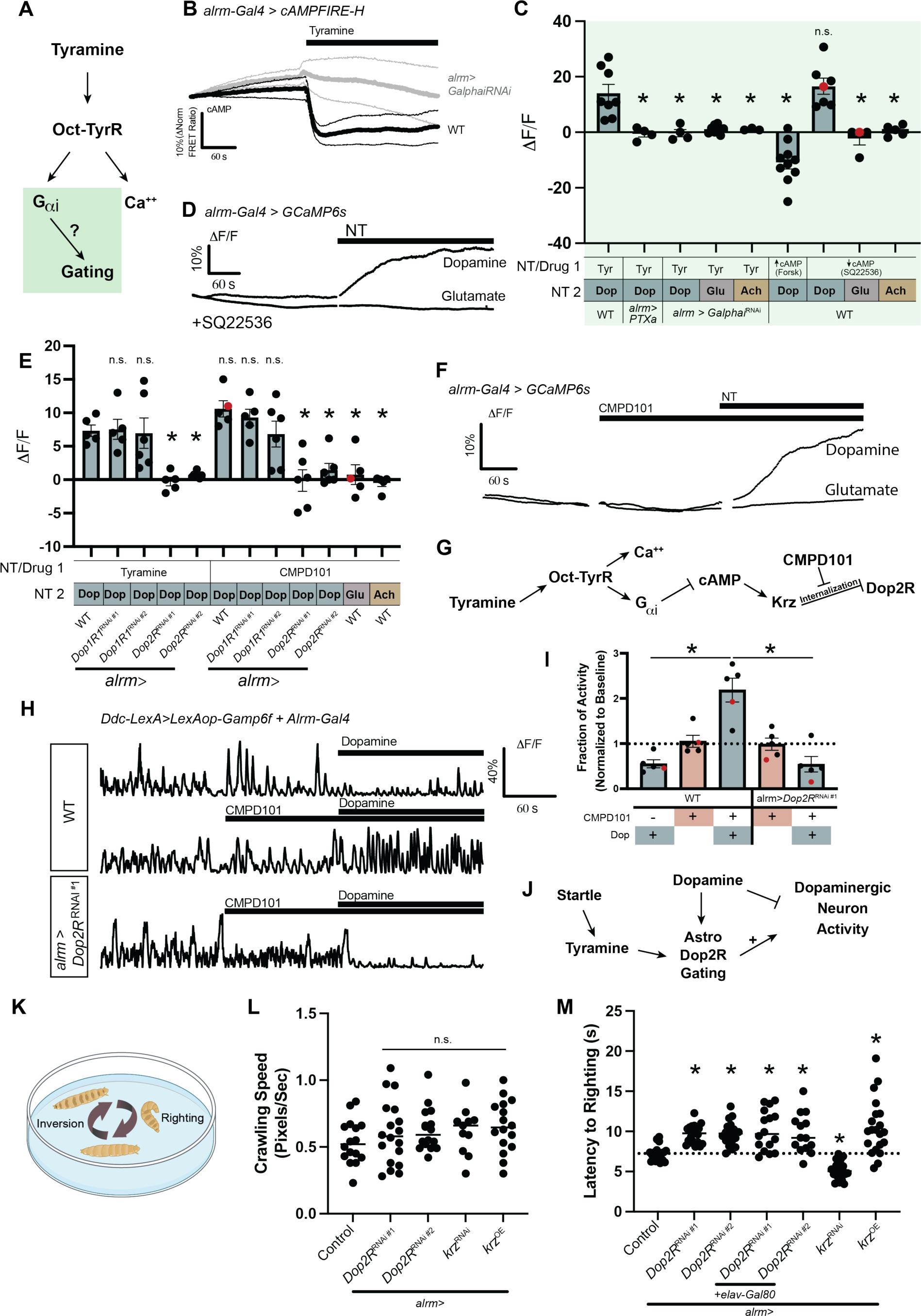
Dopamine gating is dependent on G_⍺i_ signaling. (**A**) Schematic showing hypothesized mechanism of gating via the G-protein signaling downstream of Oct-TyrR. (**B**) Mean trace (thick line) and 95% confidence interval (thin lines) of cAMP levels as measured by the FRET ratio of the cAMP indicator cAMPFIRE-H in astrocytes (n=5). Tyramine exposure causes a decrease in cAMP levels which aligns with its hypothesized function as a G_⍺i_ GPCR and this decrease is prevented by knocking down *Galphai*. (**C**) Quantification of astrocyte calcium response demonstrates that the gating of all NTs is dependent on G_⍺i_ signaling but cAMP modulation is only sufficient to gate dopamine. (**D**) Example traces of astrocytes treated with the adenylyl cyclase inhibitor SQ22536 responding to dopamine but not glutamate. (**E**) Quantification of astrocyte calcium responses shows that dopamine gating is dependent on *Dop2R* but not *Dop1R1*. Dopamine gating can also be achieved by first treating astrocytes with CMPD101 – which inhibits kinase-mediated internalization of receptors – and this gating is similarly dependent on Dop2R. (**F**) Example traces of astrocytes responding to dopamine but not glutamate following exposure to CMPD101. (**G**) Schematic showing the hypothesized mechanism that links tyramine exposure to dopamine gating. (**H**) Traces (without TTX) of *Ddc*^+^ dopamine neuron activity. Bath application of dopamine normally inhibits *Ddc*^+^ neuron activity but becomes excitatory after pre-exposure to CMPD101. Knocking down *Dop2R* in astrocytes reverts the effects of CMPD101 and bath application of dopamine becomes inhibitory once again. (**I**) Quantification of neuronal activity from experiments outlined in 2H. (**J**) Schematic showing hypothesized role of astrocyte gating of Dop2R on the response of dopaminergic neurons to bath application of dopamine (**K**) Diagram of larval righting assay. (**L**) Larval crawling analysis shows that *Dop2R* and *Krz* manipulations in astrocytes do not affect baseline larval locomotion. (**M**) Quantification of latency to right of larvae turned to their posterior side (righting). Knocking down *Dop2R* in astrocytes with or without *Gal80* expression in neurons leads to slower righting. Further, *krz* knockdown and overexpression have bidirectional effects on larval righting that align with the hypothesized role of Krz in internalizing Dop2R. * Indicates p-value < 0.05; details of statistical comparisons and exact p-values in Table S1. All error bars represent SEM. Red dots in bar graphs correspond to traces chosen as example.

What receptor could be mediating the response of astrocytes to dopamine? Two metabotropic dopamine receptors are present at the surface of larval astrocytes – one homologue of *DRD1* (*Dop1R1*) and one homologue of *DRD2* (*Dop2R*; Fig. S2). After knocking down each receptor selectively in astrocytes, we found that *Dop2R* knockdown completely prevented the response to dopamine after tyramine while *Dop1R1* knockdown had no effect (Fig. 2E). Therefore, the cAMP-mediated regulation of dopamine responses in astrocytes occurs via Dop2R.

One known mechanism by which cAMP can regulate the activity of GPCRs is via the activation of kinases that initiate internalization of receptors, which in *Drosophila* occurs via the Arrestin homologue Kurtz (Krz). We speculated that the gating of calcium responses to dopamine could occur via the inhibition of Dop2R internalization in astrocytes, permitting more receptors to remain on the surface and enhancing receptor signaling. CMPD101 is an inhibitor of kinases required for receptor internalization and we sought to determine whether it could functionally replace adenylyl cyclase inhibition in preventing receptor internalization and gating responsiveness to dopamine (Fig. 2G). Indeed, we found CMPD101 treatment successfully gated dopamine responses via Dop2R but did not affect glutamate or acetylcholine responses (Fig. 2E-F). Thus, gating of dopamine responses in astrocytes downstream of tyramine stimulation appears to occur via the cAMP-mediated modulation of Dop2R internalization (Fig. 2G). Given its lack of effect on glutamate or acetylcholine responses, our data further suggests that gating of astrocyte responsiveness to other neurotransmitters occurs via different mechanisms (i.e., not via surface retention of receptors). Protein internalization has also been proposed to underlie the astrocytic regulation of GABA levels in mammals (*22*), suggesting that surface exposure is an evolutionarily conserved mechanism to tune astrocyte sensitivity to neurotransmission.

To better understand how neurotransmitter response gating could impact information processing in the brain, we sought to identify an *in vivo* circuit specifically modulated by Dop2R signaling in astrocytes. Larval VNC astrocytes have been shown to mediate the inhibition of dopamine neurons by the arousal cue tyramine (*2*), raising the possibility that astrocyte Dop2R responses could impact state-dependent changes in dopamine circuit regulation. We expressed *GCaMP6f* in dopaminergic neurons and found that addition of dopamine in the VNC led to a reduction of dopaminergic neuron activity (Fig. 2H-I). Pretreating with CMPD101, which gates the astrocyte dopamine response, inverted this effect and caused a dramatic increase in dopaminergic neuron activity in response to dopamine application (Fig. 2H-I). Remarkably, this reversal was abolished when *Dop2R* was selectively eliminated from astrocytes, demonstrating that this inversion of dopamine’s effect was mediated by astrocyte Dop2R (Fig. 2H-I). Our data therefore reveal, unexpectedly, that Dop2R-mediated responses gated by arousal-associated tyraminergic cues in astrocytes can drive enhanced dopaminergic activity (Fig. 2J).

We next asked if the tyramine-gated dopamine response in astrocytes could influence whole-animal behavior. Rolling a larva onto its dorsal side induces a startle response that requires coordinated, dopamine-modulated motor activity to right itself (Fig. 2K) (*23*). Given the intersection of a startle response and dopamine circuit activity – and given the ability of astrocytes, which tile much of the brain, to detect dopamine input from a large array of sources – we wondered if the gating of dopamine responses in astrocytes could regulate this behavior. Knocking down *Dop2R* in astrocytes using two distinct RNAi constructs did not affect baseline larval locomotion but significantly delayed larval righting (Fig. 2L-M). This aligns with our calcium imaging of dopaminergic neurons, which suggested that the gated astrocyte dopamine response enhances dopamine circuit activity (Fig 2G). To further confirm the effect of *Dop2R* knockdown was neuron-independent, we combined astrocyte *Dop2R* knockdown with a transgene that prevents any *Dop2R* knockdown in neurons (*elav-Gal80*) and observed a similar delay in larval righting (Fig. 2M).

If the observed changes are due to arousal-mediated changes in Dop2R trafficking in astrocytes as our imaging experiments suggest, we would predict that modulation of Krz activity would also modulate larval righting. Indeed, we found that overexpressing *krz* – which should drive excess internalization of Dop2R – phenocopied *Dop2R* knockdown and slowed larval righting (Fig. 2M). Reciprocally, knocking down *krz* to retain Dop2R at the surface enhanced righting behavior (Fig. 2M). Thus, increased or decreased Krz activity can bi-directionally modulate larval righting efficiency.

Together with our results from imaging dopamine neuron activity, these findings argue that dynamic trafficking of Dop2R downstream of arousal signaling to astrocytes can powerfully modulate neuronal circuit activity with impacts at the level of whole-animal behavior.

### Astrocyte response gating by adrenergic signaling is conserved in mammals

Astrocytes can change neural circuit activity downstream of adrenergic-like signaling in many species (*2, 3, 14, 19*), arguing that the fundamental mechanisms of astrocyte circuit modulation are highly conserved. To determine if adrenergic signaling drives dopamine gating in mammalian astrocytes, we performed calcium imaging experiments in primary rat astrocytes loaded with the calcium indicator Fluo-4 (Fig. 3A-B). For each primary preparation, we first determined the highest dose of dopamine that itself could not induce a calcium response in primary astrocytes. Working at that concentration of dopamine, we found that bath application of norepinephrine (NE) was sufficient to gate the response of astrocytes to the previously subthreshold dopamine exposure (Fig. 3B-D). Thus, exposure of mammalian astrocytes to NE can also promote astrocyte responsiveness to dopamine.

**Fig 3:**
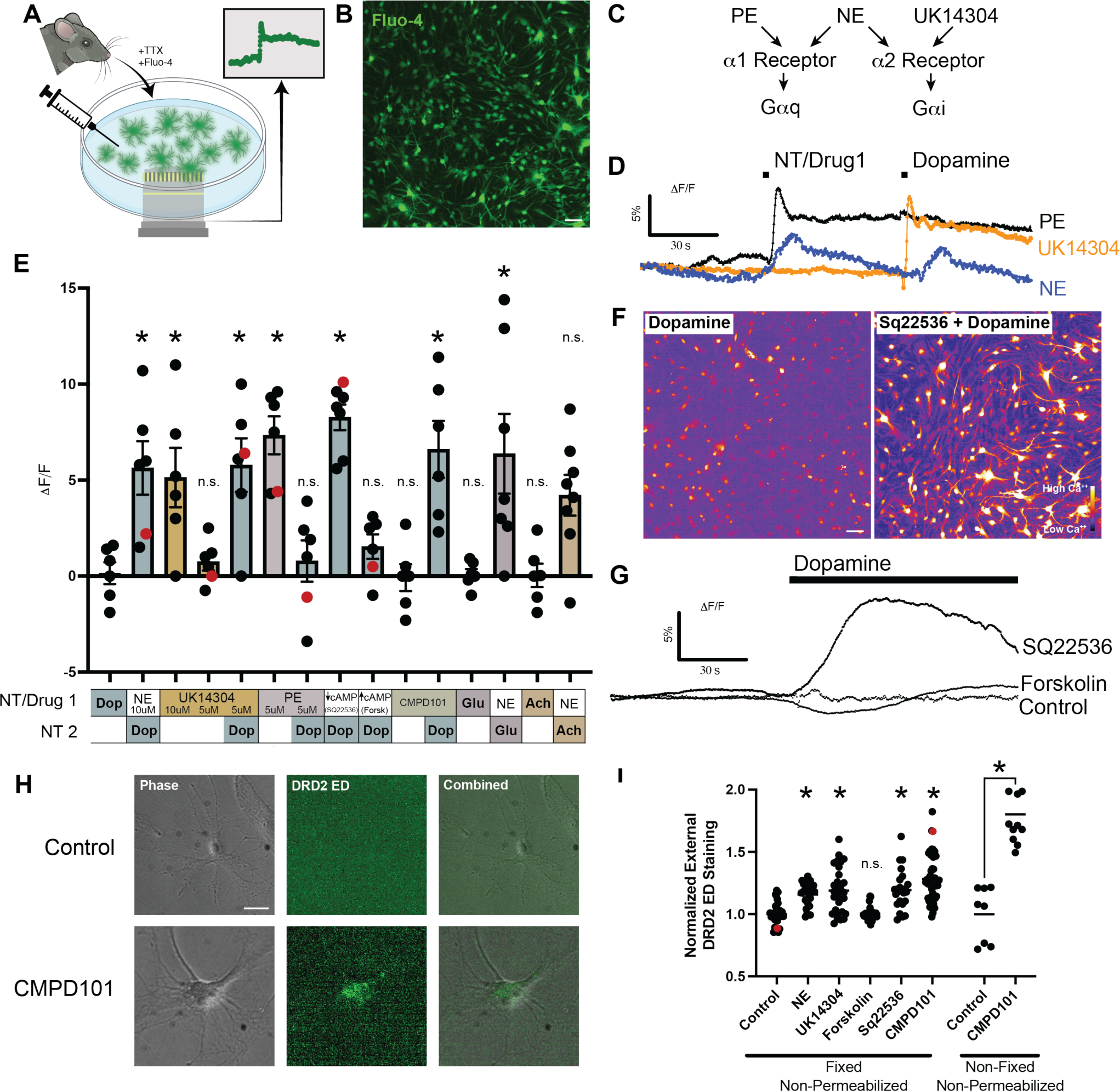
Gating of dopamine is conserved in mammals. (**A**) Schematic showing method of calcium imaging in primary rat astrocytes. (**B**) Example image of primary rat astrocytes loaded with the calcium indicator Fluo-4. (Scale bar = 20 μm) (**C**) Schematic of NE and drug selectivity for ⍺1 vs ⍺2 adrenergic receptors. (**D**) Example traces of rat astrocyte calcium responses. Both NE and the ⍺2 adrenergic agonist UK14304 can gate the response to dopamine. (**E**) Quantification of astrocyte calcium responses to various drugs and dopamine suggest that the cellular mechanism of gating is shared between *Drosophila* and mammalian astrocytes. Namely, dopamine gating can be mediated by G_⍺i_ adrenergic stimulation, cAMP modulation, and CMPD101-mediated inhibition of internalization. NE exposure can also gate glutamate and there is a trend towards NE gating acetylcholine (**F**) Example astrocyte calcium responses to SQ22536 + dopamine but not dopamine alone shown via pseudo colored Fluo-4 fluorescence. (Scale bar = 20 μm) (**G**) Example trace of rat astrocytes responding to dopamine following SQ22536 mediated inhibition of adenylyl cyclase but not to dopamine alone nor to dopamine following adenylyl cyclase activation via forskolin (**H**) Example images of astrocytes (phase) with or without treatment with CMPD101 for 30 minutes and then stained with an antibody against the extracellular domain of DRD2 (DRD2 ED; green) (Scale bar = 10 μm; brightness of the DRD2 ED staining enhanced in the example control image to demonstrate that astrocyte staining is at or near background levels). (**I**) Quantification of primary rat astrocytes stained for the extracellular domain of DRD2 shows that manipulations that can gate the dopamine response in *Drosophila* and rat also lead to increased externalization of DRD2. * Indicates p-value < 0.05; details of statistical comparisons and exact p-values in Table S1. All error bars represent SEM. Red dots correspond to traces chosen as example.

Unlike in *Drosophila* astrocytes – where calcium responses to noradrenergic homologues occur downstream of a single G_⍺i_ GPCR – mammalian astrocytes express a host of adrenergic receptors including the high affinity ⍺1 G_⍺q_-coupled and ⍺2 G_⍺i_-coupled receptors (Fig. 3C, S3). To determine if gating is mediated by G_⍺i_ signaling, we stimulated astrocytes with an ⍺1 agonist (phenylephrine, PE) or an ⍺2 agonist (UK14304) followed by a subthreshold dose of dopamine. While PE induced a robust calcium response, it was unable to gate the dopamine response (Fig. 3C-E). In contrast, while a high concentration of UK14304 could induce calcium responses, even subthreshold UK14304 stimulation was sufficient to gate dopamine (Fig. 3C-E). Thus, ⍺2 G_⍺i_ signaling in the absence of initial calcium influx is sufficient to enhance the astrocytic response to dopamine in mammalian astrocytes.

Modulation of cAMP levels and inhibition of receptor internalization were both sufficient to modulate dopamine responses in fly astrocytes. To test if these mechanisms are conserved in mammals, we modulated cAMP levels using either forskolin (↑cAMP) or SQ22536 (↓cAMP). Inhibition but not activation of adenylyl cyclase was sufficient to gate dopamine response in the absence of NE or ⍺2 agonist exposure (Fig. 3E-G). Similarly, CMPD101-mediated inhibition of receptor internalization was also able to induce responses to dopamine (Fig. 3E). Thus, the mechanism of dopamine gating via G_⍺i_ GPCR activation, inhibition of adenylyl cyclase, and inhibition of receptor internalization appears to be evolutionarily conserved from flies to mammals. Gating was also observed for glutamate following NE exposure and there was a trend towards gating of acetylcholine (Fig. 3E).

Finally, to more directly test whether our manipulations of dopamine response gating are due to changes in receptor localization, we treated astrocytes with the drugs utilized in Fig. 3E and stained using an antibody against the extracellular domain of DRD2 (Fig 3H). We quantified the amount of cell-surface DRD2 exposure by staining without permeabilization. In line with our hypothesized mechanism, exposure to NE, UK14304, SQ22536, and CMPD101 – all of which can gate dopamine responsiveness – led to an increase in DRD2 extracellular staining while forskolin – which does not lead to enhanced dopamine responsiveness – did not enhance cell surface staining (Fig. 3H-I).

## Discussion

In order to determine how astrocytes regulate neurons, it is essential to understand the mechanisms that link astrocytic neurotransmitter exposure to intracellular calcium signaling and downstream circuit modulation. Our data suggest that astrocytes dynamically alter their response to neurotransmitters based on cell state, such as whether or not the cell has recently received an arousal cue (e.g., Oct/Tyr or NE). The modulation of neurotransmitter responses downstream of NE/GPCR signaling is also conserved in mammals, arguing that it is an ancient and fundamental feature of complex nervous systems. An important question for the future will be whether the observed response enhancement during gating of mammalian astrocytes to is truly an on-or-off mechanism as appears to be the case in flies, or if it acts as more of a priming effect to increase or decrease the gain of astrocyte responses.

We also discovered that astrocyte responsiveness to different neurotransmitters is regulated differently downstream of initial GPCR signaling: dopamine is gated over long time periods by changes in cAMP and broad manipulation of receptor surface exposure, while acetylcholine and glutamate responsiveness is modulated simultaneously with tyramine exposure and does not require cAMP modulation. A longstanding mystery in neuroscience has been how a single astrocyte could specifically listen to the thousands of synapses from disparate neuronal subtypes that are contained within its territory. Local gating of each neurotransmitter response, through the signaling events we describe here, could serve as a flexible mechanism to allow astrocytes to turn on and off their sensitivity to the various circuits within their domain, or if used more broadly, to drive state-dependent changes in neuronal activity (*2, 3, 6, 24*).

At a technical level, the observation that one neurotransmitter can alter an astrocyte’s response to others suggests that differences in cell state could underpin regional and temporal variations in astrocyte responses and thus the discrepant reports of astrocyte responsiveness to neurotransmitters. For instance, the ability of glutamate to induce astrocytic calcium activity appears to vary based on brain region and age (*10-17*). Gating could potentially contribute to such differences in astrocyte responses over time and space. Recent studies have also suggested that neurotransmitter responses within one astrocyte can modulate the responsiveness of gap-junction-coupled astrocytes nearby, potentially adding variability to what different experimenters might witness in response to the same manipulation (*10*). Because gap junctions are thought to be large enough to pass cAMP, an intriguing hypothesis is that local changes in cAMP might travel between cells and mediate gating responses across the larger astrocyte syncytium.

While neuromodulation is often thought about in terms of how far a given neuromodulator could potentially diffuse, serial electron micrograph reconstructions of the brain have highlighted the dense nature of the brain parenchyma, which likely limits the long-range diffusion of neuromodulators in the extracellular space (*25*). Given the dense infiltration of astrocyte processes in the brain, the numerous ways that astrocytes can respond to neuromodulators, and the different mechanisms by which they can regulate neuronal activity, astrocytes increasingly appear as likely candidates for the long-range transmission of neuromodulatory cues through receipt and rebroadcasting of neuronal signals. Thus, astrocytic regulation of downstream neuronal activity must be considered when examining both local and broad aspects of neural circuit function, particularly in the context of slower acting neuromodulatory events.

How does gating affect specific neuronal circuits? Our work reveals that dopamine neuron activity can be potently regulated by astrocyte Dop2R signaling. Remarkably, we found that CMPD101-medidated inhibition of receptor internalization led to a two-fold increase in dopamine neuron activity via astrocytic Dop2R. This powerful bidirectional modulation of dopamine circuitry is mirrored by a bidirectional control of larval behavior: knocking out Dop2R in astrocytes regulates the speed of larval righting, a behavior that sits at the intersection of a tyramine-induce arousal signal and downstream dopamine-regulated motor actions. How astrocytes can regulate dopaminergic neuron signaling so profoundly is a crucial next question.

Our work also implies that astrocyte modulation of dopaminergic activity is complex and context-dependent. While previous studies in our lab demonstrated that calcium influx downstream of octopamine/tyramine stimulation leads to silencing of dopaminergic neurons, likely via adenosine signaling, here our data indicates that Dop2R signaling in astrocytes leads to an increase in dopamine neuron activity. We propose three alternative models to explain this diversity of signaling mechanisms: 1) the source of calcium influx (e.g., extracellular entry versus release from internal stores) or strength of calcium response can coopt different downstream signaling mechanisms and drive disparate outputs, 2) the localization of calcium influx in the cell can dictate different outputs, or 3) second messengers other than calcium that accompany metabotropic signaling are responsible for some neuromodulatory signals in astrocytes. In line with these alternative possibilities, cAMP levels can be modulated by calmodulin activity downstream of calcium influx, highlighting the complex interactions between these secondary signaling cascades. Only by continuing to decipher the mechanistic links between these various astrocyte intracellular signals (including calcium influx and GPCR signaling) and downstream neuromodulation will we likely decipher how astrocytes regulate information processing in the brain.

## Acknowledgments

We would like to acknowledge all members of the Freeman lab for thoughtful discussions and suggestions throughout this project. We acknowledge the fly community for generous sharing of reagents. We acknowledge expert technical assistance by the OHSU Advanced Light Microscopy Core (RRID:SCR_009961).

## Author contributions

Conceptualization: KAG, MRF

Investigation: KAG, IM, ES, LB, EM, LAA, RDK

Visualization: KAG, MRF

Writing – original draft: KAG, ES, LB, EM, LAA, MRF

Writing – resubmission draft: KAG, ES, LB, EM, LAA, MRF

## Data and materials availability

All data are available in the main text.

## Supplemental Figures and Tables

**Fig. S1.**
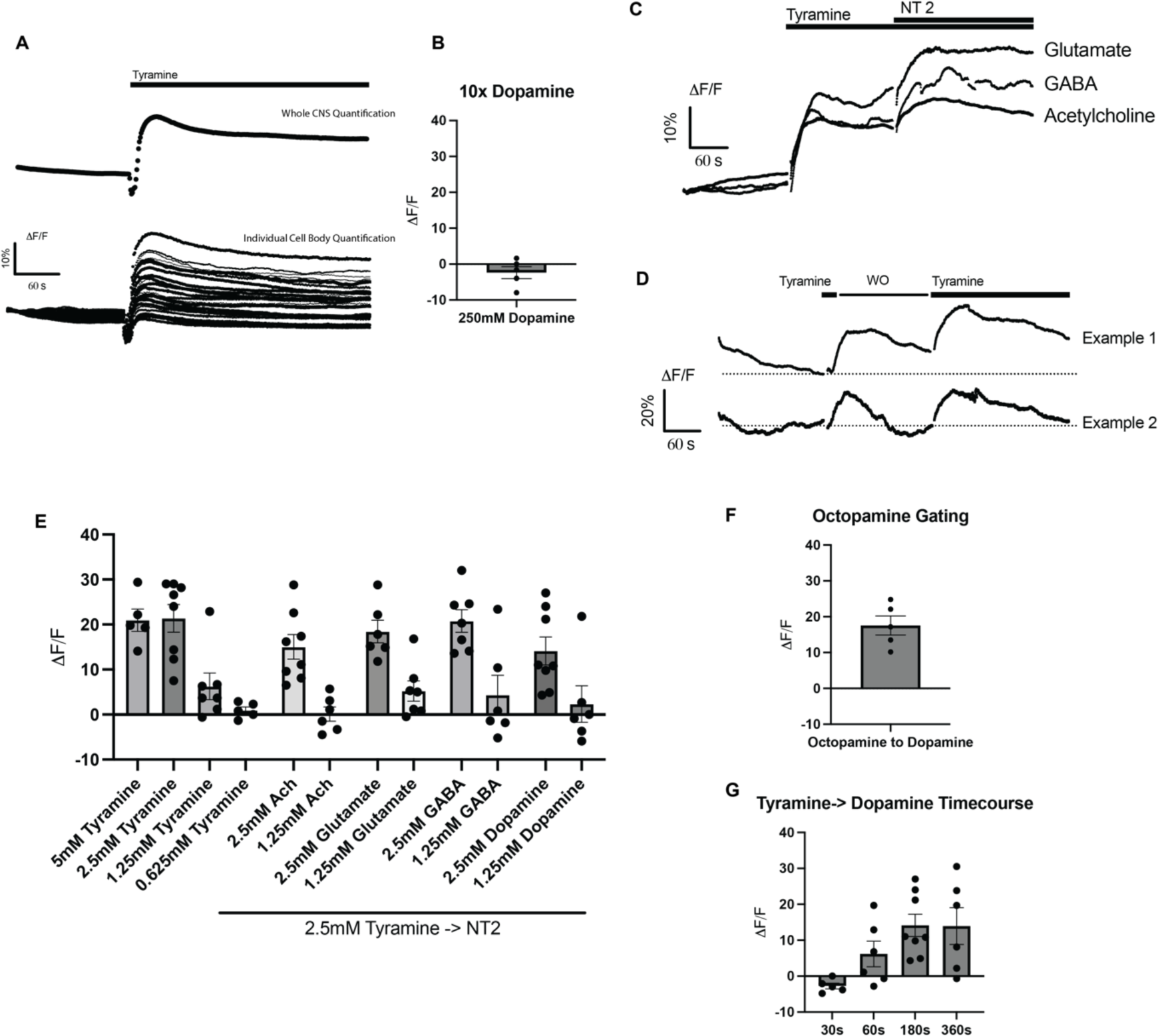
Extended sample traces of imaging and gating. (A) Comparison of quantification of whole CNS (top) vs individual astrocyte cell bodies (bottom). Note that astrocytes demonstrate a relatively homogenous response to bath application of neurotransmitter. (B) Treatment of astrocytes with 250mM dopamine (10 times the dose used in all other experiments) cannot induce a calcium response in isolation. (C) Example traces of gating responses for glutamate, GABA, and acetylcholine. (D) Example traces of tyramine responses after an initial tyramine response and washout. Note that in some animals calcium returns to baseline before subsequent neurotransmitter exposure and in some animals calcium remains elevated, but this does not obviously impact the second calcium response. (E) Varying doses of the baseline astrocyte response to tyramine as well as gating of the various neurotransmitters suggests that astrocyte responses are highly sensitive to bath neurotransmitter concentration in our *ex vivo* preparation. (F) Octopamine, which also activates the Oct-TyrR, can also gate other neurotransmitter responses. (G) Varying the time between tyramine exposure and dopamine exposure demonstrates that dopamine gating does not occur when only 30 seconds separates the two neurotransmitters and can begin to occur 60 seconds after tyramine response.

**Fig. S2.**
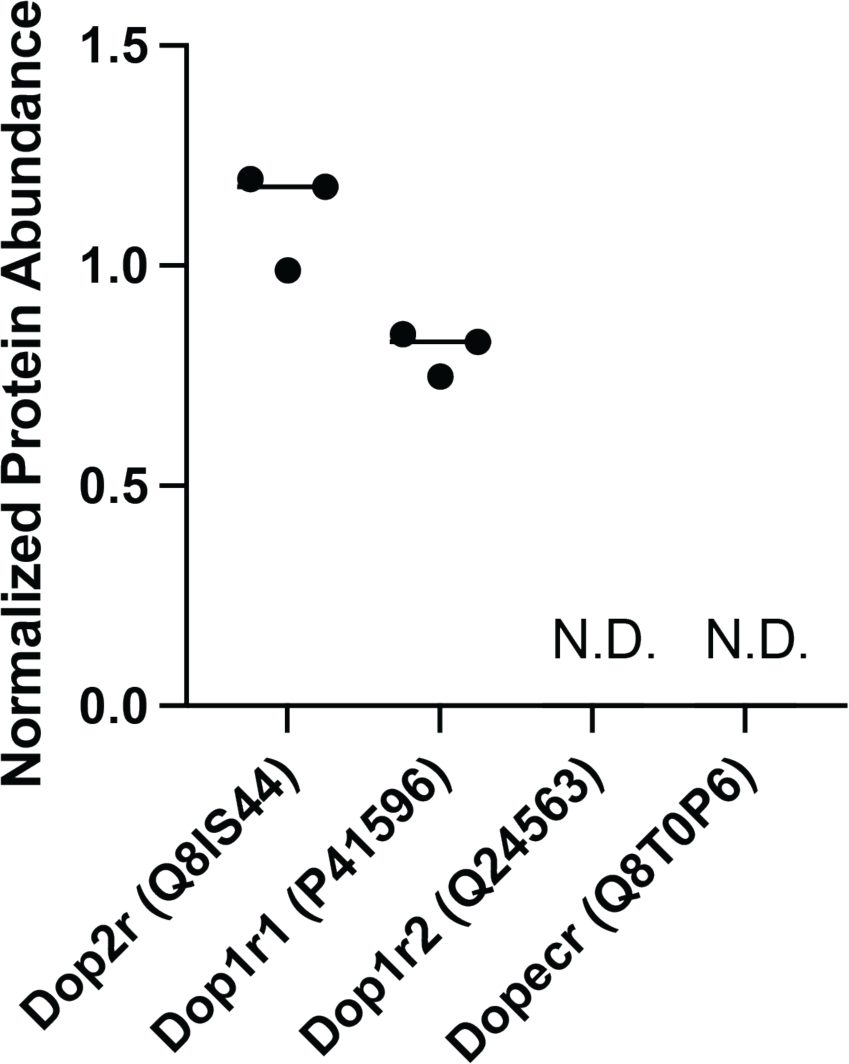
Metabotropic dopamine receptors in *Drosophila* larval astrocytes. Protein abundance of all four *Drosophila* metabotropic dopamine receptors in larval astrocytes from unpublished internal mass spectrometry dataset. N.D. indicates that no protein fragments were detected.

**Fig. S3.**
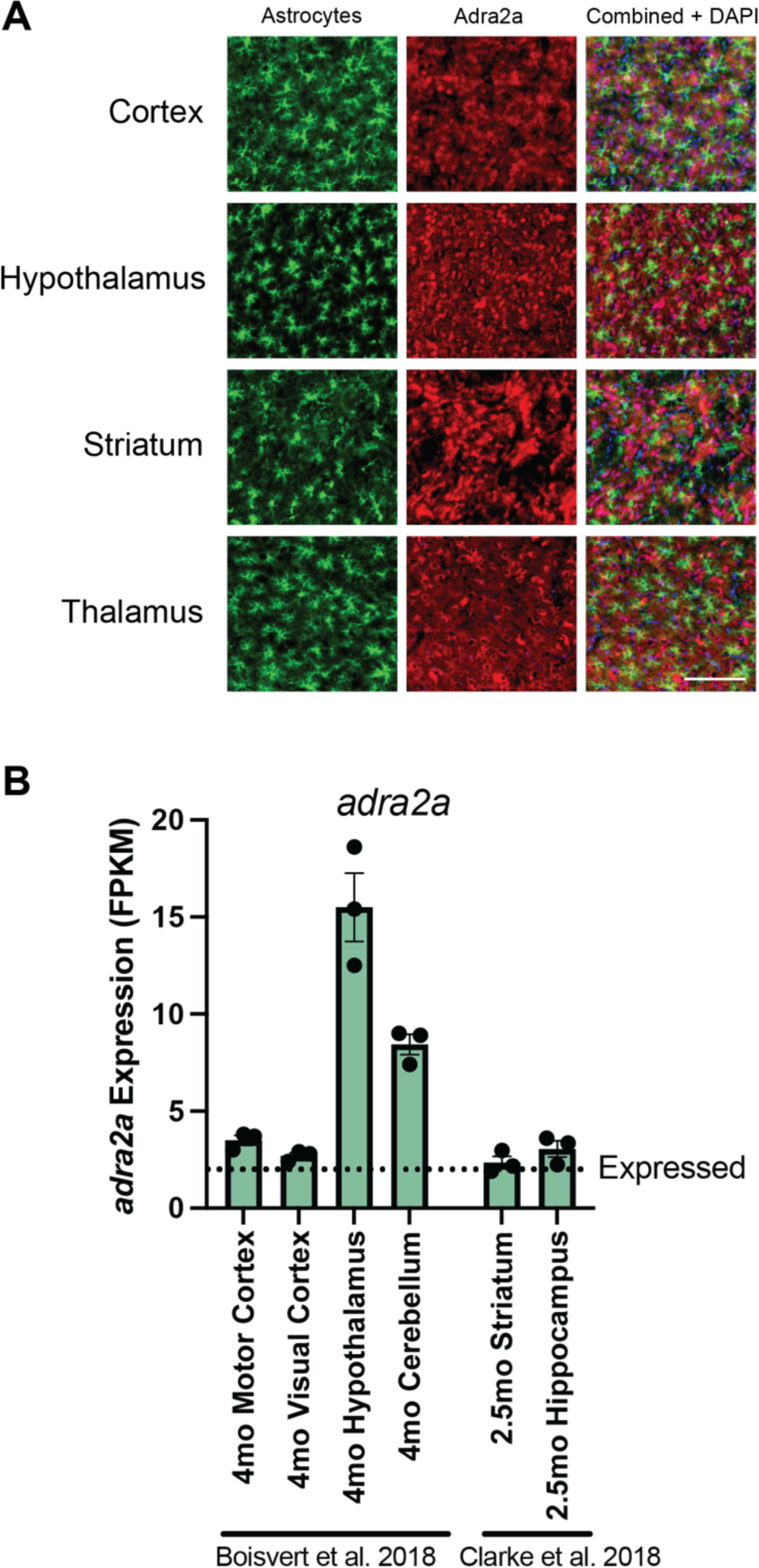
Characterization of Adra2a in the rodent brain. (A) Staining for Adra2a in various brain regions of Aldh1l1-EGFP/Rpl10a mice. Staining indicates that Adra2a is broadly present in the brain, including in astrocytes. (Scale Bar = 100 μm) (B) Previous transcriptomic databases show that mouse astrocytes isolated from various brain regions express *adra2a* (datasets from Boisvert et al., 2018 (*29*) and Clarke et al., 2018 (*30*)).

**Table S1.**
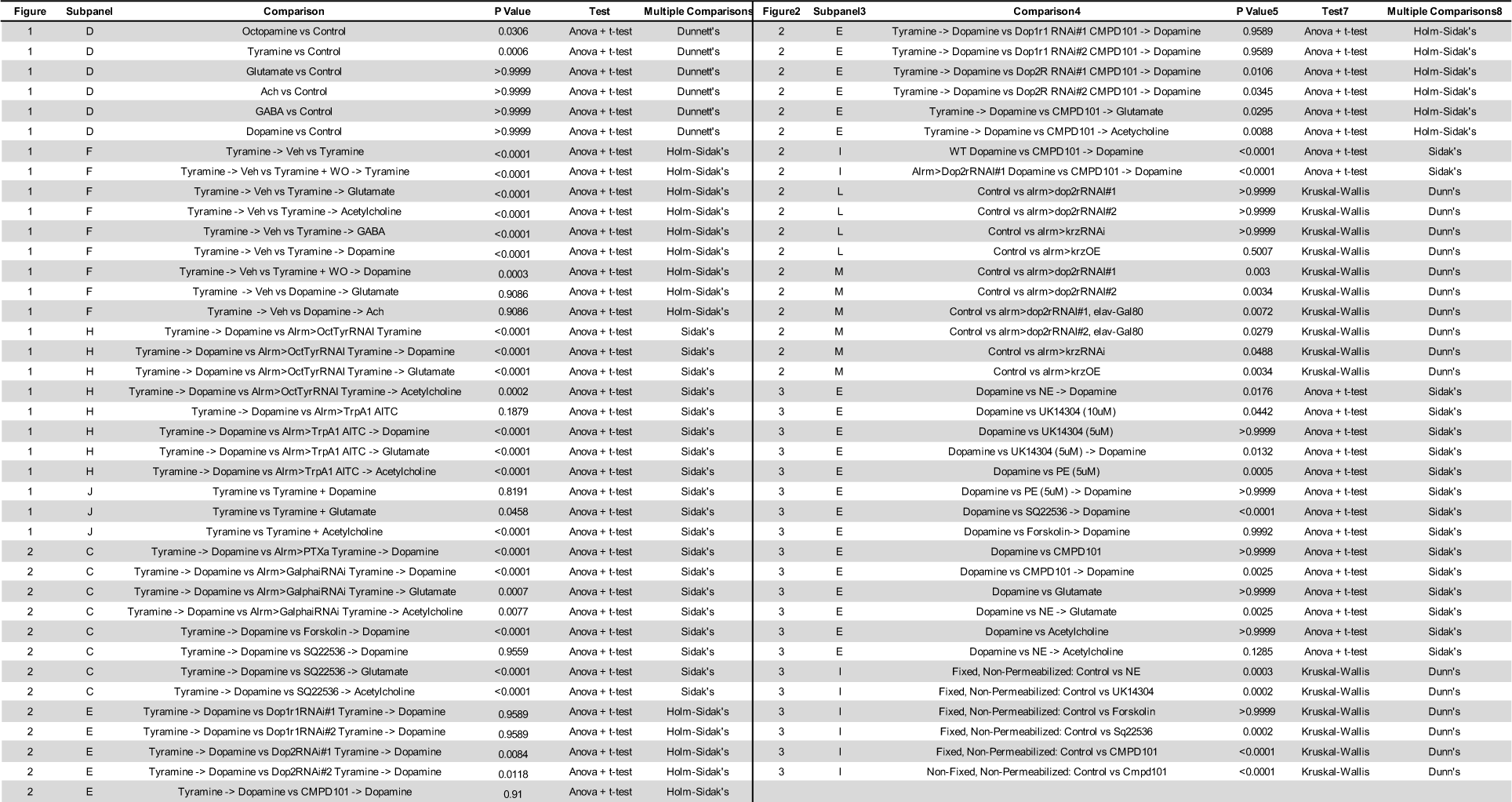
Statistical comparisons in this study.

## Materials and Methods

### Fly stocks and husbandry

Flies were kept on cornmeal food at 25 °C in 12 h/12 h light-dark cycles. Stocks utilized include: w1118 (WT/control), *alrm-Gal4* (Bloomington, 67031), *20XUAS-IVS-GCaMP6s* (Bloomington, 42746), *UAS-Oct-TyrR*^RNAi^ (Bloomington, 28332), *UAS-TrpA1* (Bloomington, 26263), *UAS-Ptxa* (Bloomington, 92006), *UAS-Galphai*^RNAi^ (Bloomington, 40890), *UAS-Dop1R1*^RNAi^ ^#1^ (Bloomington, 62193), *UAS-Dop1R1*^RNAi^ ^#2^ (Bloomington, 55239), *UAS-Dop2R*^RNAi^ ^#1^ (Bloomington, 50621), *UAS-Dop2R*^RNAi^ ^#2^ (Bloomington, 36824), *elav-Gal80* (*26*), *UAS-krz*^RNAi^ (Bloomington, 29523), *UAS-krz* (Bloomington, 27889), *Ddc-LexA* (Bloomington, 54218), *13XLexAop2-IVS-GCaMP6f* (Bloomington, 44277). *UAS-cAMPFIRE-H* were provided as a generous gift from Dr. Haining Zhong and Dr. Bing Ye (*27*).

### Drugs and neurotransmitters

All neurotransmitters in *Drosophila* imaging experiments were added at a final concentration of 2.5 mM. In primary rat astrocyte culture experiments, a dopamine concentration that was itself subthreshold to induce calcium responses was found empirically for each cell isolation and then used for all subsequent experiments – in 2/3 preparations this concentration was 2.5 mM and in one preparation this concentration was 1.25 mM. Neurotransmitters were obtained from Sigma: Glutamate (49621), Dopamine (H8502), Tyramine (T2879), Acetylcholine (A6625), GABA (A2129), Octopamine (O0250). AITC (Allyl isothiocyanate, FisherScientific AC102950050) was used at a concentration of 100 μM. Unpublished experiments in the lab suggest that higher concentrations of AITC begin to inhibit astrocyte calcium influx but not neuronal calcium influx. CMPD101 (Tocris, 5642) was used at a concentration of 1 μM. Forskolin (Tocris, 1099) was used at a concentration of 10 μM. Sq22536 (Tocris, 1435) was used at a concentration of 50 μM and *Drosophila* tissue and astrocyte cultures were exposed to the drug for 10 min to allow for sufficient inhibition of adenylyl cyclase before proceeding with experiments. Norepinephrine (NE) was acquired from Tocris (5169) and treated at a final concentration of 10 μM. UK14304 was acquired from Tocris (0425) and treated at the concentrations indicated in the figures. Phenylephrine (PE) was acquired from Tocris (2838) and treated at the concentrations indicated in the figures.

### Drosophila ex vivo imaging

Wandering 3^rd^ instar larvae were dissected in <60s using the “Larval Dissection” protocol available from Janelia Flylight Protocols while immersed in dissection media composed of 110 mM NaCl, 5.4 mM KCl, 0.3 mM CaCl2, 0.8 mM MgCl2, 10 mM d-glucose, 10 mM HEPES, pH 7.2. The dissected CNS was then transferred to a Petri dish with a base of SYLGARD-184 (Sigma, 761028) coated with Poly-D-Lysine (Sigma, A-003-E). Imaging was performed with the CNS immersed in 110 mM NaCl, 5.4 mM KCl, 1.2 mM CaCl2, 0.8 mM MgCl2, 10 mM d-glucose, 10 mM HEPES, pH 7.2. All astrocyte calcium imaging experiments were performed with 3 μM TTX (Tocris, 1078) to block neuronal activity. The only experiments performed without TTX were those in which neuronal activity was directly measured (Figure 2H-I). Samples were allowed to equilibrate in imaging media for 10 min before any experiments were performed.

GCaMP imaging was performed on an Examiner.Z1 spinning disc confocal using ZEN 2.3 (blue edition) at an image acquisition rate of 5 Hz (± shutter inaccuracy). For astrocyte quantification, a region of interest (ROI) was taken of the entire field of astrocytes and mean intensity calculated using the Mean ROI function in Zen. The mean fluorescence before neurotransmitter addition was calculated by taking the simple average of the 100 frames before addition. For initial neurotransmitter addition this 100 frames occurred before any neurotransmitter was added and for the second neurotransmitter a new baseline was calculated from the last 100 frames of the previous neurotransmitter addition. The peak of calcium response was then selected manually blind to condition and genotype by selecting the highest value of the initial response to neurotransmitter after neurotransmitter addition. This process was performed manually because the time to peak fluorescence varied and because manual selection prevented any fluctuations in fluorescence level that occur after the original response from biasing the results (see fluctuation in fluorescence level after initial response in Figure 1E). The ratio of the calcium response peak to the average baseline was then calculated as a percent. Example traces were normalized to the first 50 frames of intensity. For neuronal quantification, ROIs were taken of 4-8 regions of dopamine neurites and their mean intensities calculated in ImageJ. Each region was normalized to baseline and a moving minimum function of +/- 20 frames used to correct any drift in baseline. Activity was then quantified as the integral of the average baseline-corrected mean intensities of all ROIs for the 200 frames before adding the first drug (baseline), the 200 frames after the first drug was added, and the 200 frames after adding the second drug. The ratio of activity was then calculated by simple normalization of the post-drug activities to the baseline activity.

cAMP imaging was performed on a Zeiss LSM 980 using Zen (blue edition). A region of interest (ROI) was taken of the entire field of astrocytes and FRET (fluorescence energy resonance transfer) signal taken as a simple ratio. The mean FRET ratio before and after stimulation was then selected manually and the ratio used to calculate ΔF/F as a percent.

Of note, for *ex vivo* imaging of astrocytes using this method: Astrocytes are sensitive to the calcium buffering caused by GCaMP constructs. Thus, crosses of the *alrm-gal4* and *UAS-GCaMP* lines should be regularly recrossed from parent stocks, as fly stocks with persistent astrocyte-driven GCaMP expression appear to accumulate mutations that suppress GCaMP activity. Eliciting consistent calcium responses in astrocytes is also highly dependent on the age of the larvae and great care should be taken to obtain true wandering 3^rd^ instar larvae. We have also found (unpublished) that newer generations of GCaMP (GCaMP7 and GCaMP8) are less consistent than GCaMP6 in visualizing responses, potentially due to the higher apparent resting level of calcium in astrocytes vs neurons. We have also found myristoylated GCaMP constructs to show higher sensitivity to neurotransmitter gating than cytoplasmic constructs, but cytoplasmic constructs were utilized in this study due to lower levels of artifacts from drug administration compared to myristoylated constructs.

All *Drosophila* experiments were performed on at least two independent crosses of the indicated genotype and were replicated weeks to months apart.

### Larval behavior analysis

Wandering 3^rd^ instar larvae were lightly rinsed with water to remove any food on their body and then placed on a grape plate and allowed to acclimate for 1 min. The larvae were then observed and rolled until their ventral side was up using a paintbrush during a bout of forward locomotion to ensure that the larvae were not in a state of rest when rolled. The latency to righting was then quantified as the time from the onset of head movement to first forward progress on ventral side (*23*). All righting experiments were performed blind to genotype. All experiments were performed from at least two independent crosses of the indicated genotype and were replicated weeks to months apart.

Larval crawling behavior was assessed via the frustrated total internal reflection imaging method and analyzed using FIMTrack software (*28*). 4-5 larvae of the same genotype were placed on a 0.8% agar surface positioned above an IR camera after brief rinsing in PBS. Larval position was recorded for 2 min and crawling paths for each larva were automatically generated by FIMTrack. Information of average crawling rate was exported to GraphPad Prism for analysis.

### Primary rat astrocyte culture

All animal procedures were conducted in accordance with guidelines from the National Institutes of Health (NIH) and OHSU’s IACUC. P6 Sprague Dawley rats were acquired from Charles River (Strain 400). Astrocytes were purified by magnetic-activated cell sorting (MACS) from P6 Sprague Dawley rat forebrains using the Astrocyte Isolation Starter Kit, rat < P7 from Miltenyi Biotec (130-096-052). In short, rat forebrains were dissected in DPBS (Gibco, 14190) by blunt dissection including removal of meninges. Tissue was then cut into ∼1mm^3^ pieces which were enzymatically dissociated and triturated according to the manufacturer’s protocol. Debris (primarily myelin debris) was then removed using Debris Removal Solution (Miltenyi Biotec, 130-109-398) and the resulting cells were magnetically sorted for GLAST^+^ astrocytes according to the manufacturer’s protocol (Miltenyi Biotec, 130-096-052).

Astrocyte-enriched cultures were then plated on PDL-coated tissue culture plates in serum-containing astrocyte media composed of DMEM, high glucose (Gibco 11965), 10% heat-inactivated fetal bovine serum (FisherScientific, FB12999102), 50 units/ml Penicillin/Streptomycin (ThermoFisher, 15070063), and 2mM L-glutamine (FisherScientific, 25-030-081) and cultured at 37 °C and 10% CO2. After 2 days, astrocytes were switched to AstroMACS Medium (Miltenyi Biotec, 130-117-031) and cultured for 3-5 more days before imaging.

To image, astrocytes were first washed 2x with 37 °C PBS to remove dye-containing media. Cells were then loaded with the calcium indicator Fluo-4 with AM tail (1μM, ThermoFisher, F14201) for 5 min in PBS before washing 2x with primary astrocyte imaging media (NaCl 140 mM, KCl 5 mM, CaCl2 2 mM, MgCl2 2 mM, glucose 20 mM, and HEPES 10 mM, pH 7.4). Several other astrocyte primary imaging media compositions showed no calcium responses to any NTs tested but showed dramatic calcium responses to buffer-only fluid flow. All primary astrocyte imaging experiments were performed with 3 μM TTX to block any activity of possible contaminating neurons in the culture.

Live cell imaging was performed using an inverted Nikon TiE microscope equipped with a Yokogawa CSU-W1 spinning disk confocal head, running NIS Elements software (4.51.01). The cells were maintained at 10% CO2 and 34.5 °C. Images were acquired with a Zyla v5.5 sCMOS camera (Andor) with a S Plan Fluor ELWD 20x (NA 0.45) objective. ROIs were selected using the entire field of imaging and mean intensity calculated in the time measurement function of NIS-Elements. ΔF/F as a percent was then calculated manually. Individual data points in primary culture analysis represent average response of all cells in a field of view from one well and all results represent data points from at least 3 different independent primary cell collections that occurred over many months.

To assess DRD2 externalization in fixed but non-permeabilized primary rat astrocytes, we took astrocytes treated with the indicated drugs for 30 minutes (see concentrations above) and fixed the cells without permeabilization using 1% ice cold PFA for 1 minute. We blocked the cells for 1 hr at room temperature using 10% fetal bovine serum (FisherScientific, FB12999102) in PBS. We then incubated the cells overnight at 4 °C in PBS with 1% fetal bovine serum and 1:100 rabbit polyclonal antibody to the extracellular domain of DRD2 (alamone labs, ADR-002). After washing, cells were incubated for 2 hrs at room temperature in 1% fetal bovine serum in PBS with 1:250 Alexa Fluor® 488 AffiniPure™ Donkey Anti-Rabbit IgG (H+L) (Jackson ImmunoResearch, 711-545-152; RRID: AB_2313584). Cells were then washed and imaged in PBS on the inverted Nikon TiE microscope equipped with a Yokogawa CSU-W1 spinning disk confocal head. Cells were chosen at random for imaging by phase and then imaged in both phase and 488 channels using identical imaging conditions. Individual points represent averages of all cells from a well, and data was obtained from at least 3 different primary cell preparations that occurred over several months.

To assess DRD2 externalization in non-fixed, non-permeabilized primary rat astrocytes, we treated astrocytes with 1μM CMPD101 for 30 minutes or left them untreated. We then incubated them in PBS with a 1:100 dilution of the extracellular DRD2 antibody. We washed the cells gently with PBS and then incubated for 30 minutes in a 1:250 dilution of Alexa Fluor® 488 AffiniPure™ Donkey Anti-Rabbit IgG (H+L) in PBS. Cells were then washed and imaged in PBS. Cells were chosen at random for imaging by phase and then imaged in both phase and 488 channels using identical imaging conditions. Individual points represent averages of all cells from a well and data was obtained from at least 3 different primary cell preparations that occurred over several months.

### Mouse tissue immunohistochemistry

All animal experiments were approved by NYU Grossman School of Medicine’s institutional Animal Care and Use Committee. For antibody staining, Aldh1l1-EGFP/Rpl10a (RRID: IMSR_JAX:00064) mice were used. Mice were housed on a 12h light/dark cycle at 22-25 °C and 50-60% humidity and were given food and water ad libitum.

Mice were euthanized with CO2 and brains were collected following transcardiac perfusion with cold phosphate buffered saline (1x PBS) pH 7.4, followed by freshly made 4% paraformaldehyde (PFA) in PBS. Following overnight incubation in 4% PFA in PBS at 4 °C, brains were transferred to 30% w/v sucrose in PBS and stored at 4 °C. Brains were coronally sectioned using a cryostat at a thickness of 40 μm.

Free-floating 40 μm coronal brain sections were washed in 1x PBS, then blocked with 5% normal goat serum (Thermo Fisher Scientific 16210064) in 0.3% Triton X-100 in PBS (0.3% PBST) for an hour, followed by an overnight incubation in 2.5% normal goat serum in 0.1% PBST with polyclonal chicken anti-GFP (1:1000; abcam ab13970) and polyclonal rabbit anti-ADRA2a (1:50; antibodies.com A11568) at room temperature on a shaker. Following primary antibody incubation, sections were washed with 1x PBS and incubated for 1 h in 2.5% normal goat serum in 1x PBS with goat anti-chicken Alexa fluor 488 (1:500; Invitrogen A-11039) and goat anti-rabbit Alexa fluor 647 (1:500; Invitrogen A-21245). Sections were washed with 1x PBS, counterstained with DAPI (Invitrogen D1306), mounted onto slides, and coverslipped with Fluoromount-G (SouthernBiotech 0100-01). Sections were then imaged using a Keyence BZ-X710.

